# Hybrid Solid-Liquid Optics Enable Scalable, High-Resolution, Multi-Immersion Light-Sheet Microscopy

**DOI:** 10.1101/2025.11.02.685475

**Authors:** Cheng Gong, Pauline Affatato, Matt Fay, Sudha R. Guttikonda, Nathan J. O’Connor, Emerson Noble, Maci Heal, Benjamin Haydock, Renee Mapa, Estanislao Daniel De La Cruz, Johanna E. Kowalko, Maria Antonietta Tosches, Charles R. Gerfen, Rene Hen, Christopher D. Makinson, Hanina Hibshoosh, Jack Glaser, Raju Tomer

**Author notes:** Arnas Technologies, Essex Junction, VT 05452, USA.

## Abstract

Modern biology increasingly depends on data-driven discovery, requiring scalable and affordable high-content 3D imaging across molecular to organ scales. Although tissue clearing, expansion microscopy, and light-sheet microscopy (LSM) enable subcellular-resolution imaging of intact specimens, their scalability remains fundamentally limited by detection optics: immersion objectives deliver high-resolution, aberration-free imaging but with short working distances, high cost, and multi-immersion incompatibility, while air objectives offer long working distances and portability at lower cost but suffer from severe aberrations and reduced photon collection when imaging immersed samples. We introduce the Hybrid Solid-Liquid Immersion Lens (HySIL) framework, which pairs an off-the-shelf solid optical component with a refractive index-matched liquid to precompensate aberrations and enhance resolution. Building on HySIL, we developed SCOPE and Super-SCOPE, objective-agnostic imaging devices achieving submicron lateral resolution (<0.75 µm) across centimeter-scale samples using inexpensive air objectives with >30 mm working distances. Integration with a low-cost LSM platform yielded a compact, scalable system demonstrated for multi-immersion, multi-color, subcellular-resolution mapping of cleared or expanded mouse, salamander, and cavefish brains, human iPSC-derived organoids, and 3D histopathology of breast tissue. HySIL and SCOPE establish an accessible foundation for scalable, high-resolution volumetric imaging, advancing data-driven biological discovery.

## INTRODUCTION

Breakthroughs such as AlphaFold and ESM3 for protein structure prediction^1,2^, advances in computational histopathology^3^, and generative modeling of virtual cells^4^ exemplify the transformative power of data-driven approaches in life sciences. Extending this paradigm from molecular to multicellular systems require imaging technologies capable of routinely generating subcellular-resolution volumetric datasets from intact tissues and organs. Recent developments in tissue clearing and expansion microscopy (ExM)^5,6^ have provided unprecedented optical access to intact specimens. In parallel, light-sheet microscopy (LSM) has emerged as the leading modality for high-speed volumetric imaging, offering minimal phototoxicity and exceptional optical efficiency^7^. Open-source initiatives such as openSPIM^8^ and mesoSPIM^9,10^ have broadened access to LSM, while our open-source projected light-sheet microscopy (pLSM) platform^11^ has further reduced optical and mechanical footprints and material costs by orders of magnitude through repurposing consumer-grade components.

However, achieving scalability in imaging volume, sample throughput, and widespread adoption while maintaining subcellular resolution remains constrained by fundamental limitations in detection optics. Immersion objectives enable aberration-free, high numerical aperture (NA) detection necessary for achieving submicron resolution but also impose severe constraints: short working distances limiting imaging depth, high costs, narrow refractive index (RI) correction ranges limiting multi-immersion compatibility, and operational complexity requiring sealing within imaging chambers. Air objectives offer long working distances, cost reductions of several orders of magnitude, and mechanical simplicity for integration into low-cost, portable platforms. However, they suffer from severe spherical aberrations, reduced photon collection, and degraded resolution when imaging immersed samples, limiting performance to cellular resolution. Existing correction approaches including adaptive optics^12^, custom meniscus lenses^13^, and specialized corrected air objectives^14^, mitigate some of these effects but remain system-specific, require custom engineering, or are expensive. Consequently, current LSM systems often sacrifice imaging performance for portability, performing adequately for cellular-resolution applications but leaving efficient subcellular-resolution imaging of large volumes inaccessible. This limitation has become particularly pressing with the advent of ExM, which can increase specimen dimensions by an order of magnitude, creating urgent demand for aberration-free imaging over extended working distances.

To address this challenge, we developed the Hybrid Solid-Liquid Immersion Lens (HySIL) framework that generalizes the concept of the solid immersion lens (SIL)^15–17^, which was originally introduced to enhance imaging resolution by positioning a high-RI hemispherical solid element in contact with specimens, thereby increasing effective detection NA by *n* (the refractive index of the lens). While SILs have proven valuable in semiconductor photoluminescence^18^ and super-resolution microscopy^19^ applications, their use has been limited to surface imaging^19^. The HySIL approach, instead, combines a simple, off-the-shelf solid lens element with a precisely RI-matched liquid to form a continuous solid-liquid hybrid optical system. This architecture establishes two complementary optical environments: a solid interface enabling use of air detection optics, and a liquid interface allowing flexible sample positioning and imaging.

Building on this concept, we developed SCOPE (Solid-Liquid hybrid immersion-Corrected Optics for Phase Errors), an objective-agnostic modular imaging device, and its variant Super-SCOPE, which utilizes Weierstrass super-hemispherical geometry^16,17^. These devices: (a) increase effective detection NA (scaled by *n* and *n*^2^ for SCOPE and Super-SCOPE, respectively), (b) correct spherical aberrations, (c) provide tolerance to diverse immersion media, and (d) preserve the long working distance, chromatic correction, low cost, and simplicity of air objectives.

We validated SCOPE through optical simulations and point spread function (PSF) measurements. These analyses demonstrated aberration-free, submicron resolution (<0.75 µm) imaging using inexpensive air objectives with long working distances (~34 mm) that maintain high performance across diverse tissue clearing media. We then integrated SCOPE with our low-cost pLSM-based LSM platform to create a compact, cost-effective, and scalable 3D imaging system for submicron resolution imaging of centimeter-scale samples. SCOPE enabled imaging of diverse specimens, including expanded mouse and salamander brains, CUBIC-R cleared intact mouse brains, iDISCO cleared cavefish brains, brain organoids with microglia, and 3D histopathology of human breast tissue. In all cases, SCOPE produced high-contrast, color-corrected, aberration-free imaging throughout entire sample volumes. Altogether, SCOPE provides a simple, low-cost, and scalable solution that transforms air objectives into high-resolution, aberration-free, centimeter-scale detectors, bridging affordability and performance for next-generation, data-driven biological discovery.

## RESULTS

### Design and Theoretical Validation of SCOPE and Super-SCOPE

Achieving high-resolution, large-volume imaging with inexpensive, long-working-distance air objectives requires overcoming the spherical aberrations, resolution loss, and signal-collection inefficiency that arise when imaging through high-RI media typical of cleared tissues (**Fig. 1a-b**). To address these challenges, we developed the HySIL framework (**Fig. 1c-d**), which consists of a solid truncated spherical lens coupled with a precisely RI-matched immersion oil to form a continuous hemispherical optical interface. Similar to a classical hemispherical SIL (**Fig. 1c**), HySIL precompensates phase errors introduced by high-RI specimens and increases effective detection NA (**Fig. 1d**). However, unlike conventional SILs, which are inherently limited to surface-proximal imaging^19^ (**Fig. 1c**), HySIL decouples the solid lens from both the sample and the objective, enabling aberration-free imaging throughout large, cleared specimens.

**Figure 1.**
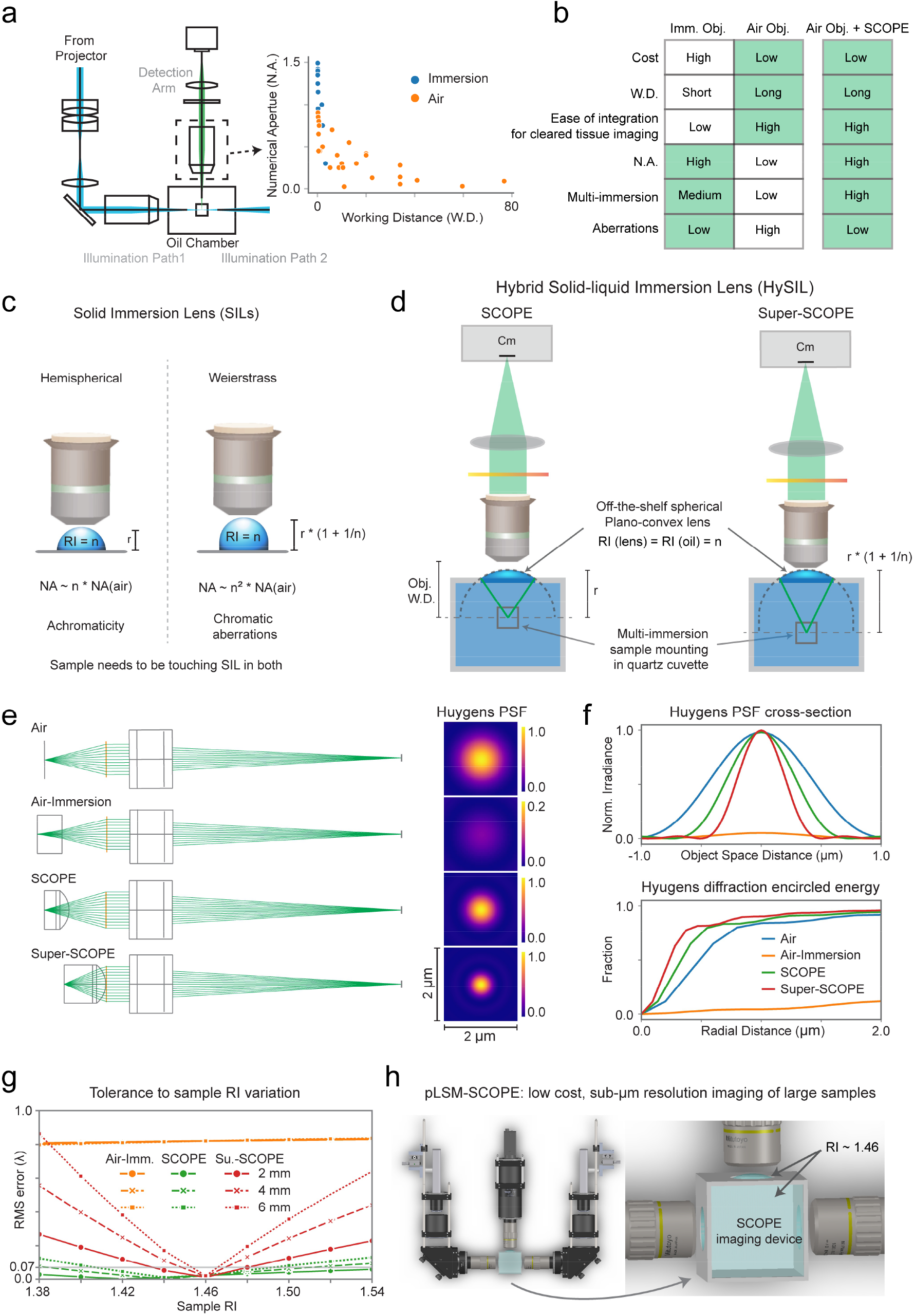
Design and validation of the HySIL and its imaging device implementations, SCOPE and Super-SCOPE. **(a)** Schematic of the projected light sheet microscopy (pLSM) setup illustrating the use of air versus immersion detection objectives. The accompanying plot compares numerical aperture (NA) and working distance (WD) for air (orange) and immersion (blue) objectives, highlighting the trade-off between resolution and scalability. Note that Illumination path 2 (not shown) mirrors path 1. **(b)** Comparative summary of key performance characteristics of immersion, air, and SCOPE-enabled air objectives. **(c)** Conceptual illustration of classical hemispherical and Weierstrass solid immersion lenses (SILs), showing NA scaling, chromatic properties, and limitations to surface-proximal imaging. **(d)** Schematic of the HySIL concept and its implementation as the modular SCOPE and Super-SCOPE imaging devices. A truncated hemispherical quartz lens is paired with refractive-index (RI)–matched immersion oil (RI ~ 1.46 in this study) to form a hybrid solid-liquid optical system that corrects aberrations while maintaining the long WD, ease of integration, and low cost of air objectives. **(e)** Zemax optical simulations comparing detection performance for air-objective imaging in air, immersion media, SCOPE, and Super-SCOPE configurations. Corresponding estimated Huygens point-spread functions (PSFs) are shown on the right. See also **Supplementary Figure 1a** for Zemax simulations of SCOPE configuration experimentally used in this study. **(f)** Quantification of normalized irradiance and diffraction-encircled energy for the Huygens PSFs shown in (e). **(g)** Tolerance analysis of air-objective detection in immersion, SCOPE, and Super-SCOPE configurations. Root-mean-square (RMS) wavefront error is plotted as a function of sample RI and thickness. **(h)** Computer-aided design (CAD) model demonstrates modular integration of the SCOPE imaging device with the compact, low-cost pLSM platform.

To translate the HySIL concept into practical platforms for enhanced light-sheet microscopy, we designed SCOPE and Super-SCOPE modular imaging devices (**Fig. 1d**), which can be seamlessly integrated into most existing LSM implementations. SCOPE incorporates an off-the-shelf plano-convex quartz lens (RI ~ 1.46) paired with a precisely RI-matched immersion oil (RI ~ 1.46) to form a hybrid solid-liquid immersion system with hemispherical geometry that increases effective NA by *n*. Super-SCOPE utilizes Weierstrass super-hemispherical geometry to achieve NA scaling by *n*^2^, providing enhanced resolution, though with reduced tolerance to optical variations. The RI of ~ 1.46 was specifically selected to ensure broad compatibility with commonly used tissue clearing methods, from aqueous (RI ~ 1.33) to solvent-based (RI ~ 1.56) protocols, without requiring modifications. The chamber geometry is calibrated such that the distance between the light-sheet illumination plane, entering through the center of the illumination window (either flat or HySIL-enabled for thinner light sheets), and the curved surface of the solid lens equals the lens radius of curvature (r) or r*\**(1*+*1/*n*) (**Fig. 1d**), ensuring optimal wavefront correction and alignment of the illumination and detection planes. Note that although we chose a truncated hemispherical plano-convex lens as the solid component for its simplicity and easy, low-cost, off-the-shelf availability in various shapes and sizes, the flexibility of the HySIL framework in using RI-matched solid and liquid components allows for the use of a wide range of other lens geometries.

We performed Zemax optical simulations to evaluate the performance of SCOPE and Super-SCOPE for high-resolution imaging of large samples cleared with diverse tissue-clearing methods (**Fig. 1e-g**). The detection system was modeled with an idealized optical path comprising a 40 mm focal length detection lens (NA_air = 0.28, matching the air objective used in this work) and a 200 mm tube lens (TTL200, Thorlabs). For direct comparison of performance across four configurations, including air, immersion, SCOPE, and Super SCOPE, a hemispherical curvature of 16 mm was selected. To emulate the configuration used experimentally to utilize most of the detection working distance, a radius of curvature of 27.5 mm was also simulated (**Supplementary Fig. 1a-c**), yielding similar results.

As shown in **Fig. 1e-g** and **Supplementary Fig. 1a**, the HySIL-based SCOPE and Super-SCOPE optical designs markedly reduced spherical aberrations, achieving diffraction-limited performance (**Fig. 1e-f**) and resolution enhancement approximately by factors of *n* and *n*^2^, respectively (compare PSFs in **Fig. 1e-f**). Wavefront error analysis further confirmed effective compensation of optical path differences across both sagittal and tangential planes (**Supplementary Fig. 1b**), demonstrating efficient precompensation of phase errors introduced when imaging samples immersed in high-RI media.

We next assessed the tolerance of SCOPE and Super-SCOPE (system RI ~ 1.46) across a range of sample RIs representative of common tissue clearing protocols (1.38 to 1.54) and sample thicknesses of 2 to 6 mm. As shown in **Fig. 1g** and **Supplementary Fig. 1c**, the root mean square (RMS) wavefront error remained consistently below or near the Maréchal criterion (λ/14) under all tested conditions for SCOPE, confirming high correction efficiency and broad adaptability without hardware adjustments. As expected, Super-SCOPE exhibited more limited tolerance to RI variation but still outperformed conventional air objective-based imaging of immersed samples.

Collectively, these optical simulations establish HySIL as a generalizable framework for aberration-free, high-resolution imaging of large, cleared samples using inexpensive, long-working-distance air objectives. The passive, easy-to-integrate design of SCOPE and Super-SCOPE chambers eliminates the need for sealed immersion assemblies, allowing seamless integration into a wide range of imaging systems, including our lab-built pLSM and its commercially available implementation, the SLICE (MBF Bioscience) platform.

### Experimental Validation of SCOPE through PSF Characterization

To experimentally validate SCOPE performance, we measured and characterized the PSF of the detection optics using two inexpensive long-working-distance air objectives: Mitutoyo 10x/0.28 NA/34 mm WD and 5x/0.14 NA/34 mm WD. The SCOPE imaging device incorporated a plano-convex fused quartz lens (RI ~ 1.46; radius of curvature of 27.5mm) paired with an RI-matched immersion oil to form the hybrid solid-liquid immersion system.

We first imaged sub-resolution (150 nm) gold nanoparticles embedded in 0.5% agarose (H_2_O) under wide-field illumination to assess the wide-field detection PSF of SCOPE (**Fig. 2a-b**). Identical illumination power was applied for both SCOPE and conventional air-immersion setups to enable direct comparison of resolution and signal collection efficiency. As shown in **Fig. 2a-b**, SCOPE effectively corrected spherical aberrations across multi-millimeter scale fields of view (1.2 mm for 10x and 2.7 mm for 5x objectives) and produced higher signal intensity relative to the Air-Immersion configuration (compare lookup tables in **Fig. 2a-b**).

**Figure 2.**
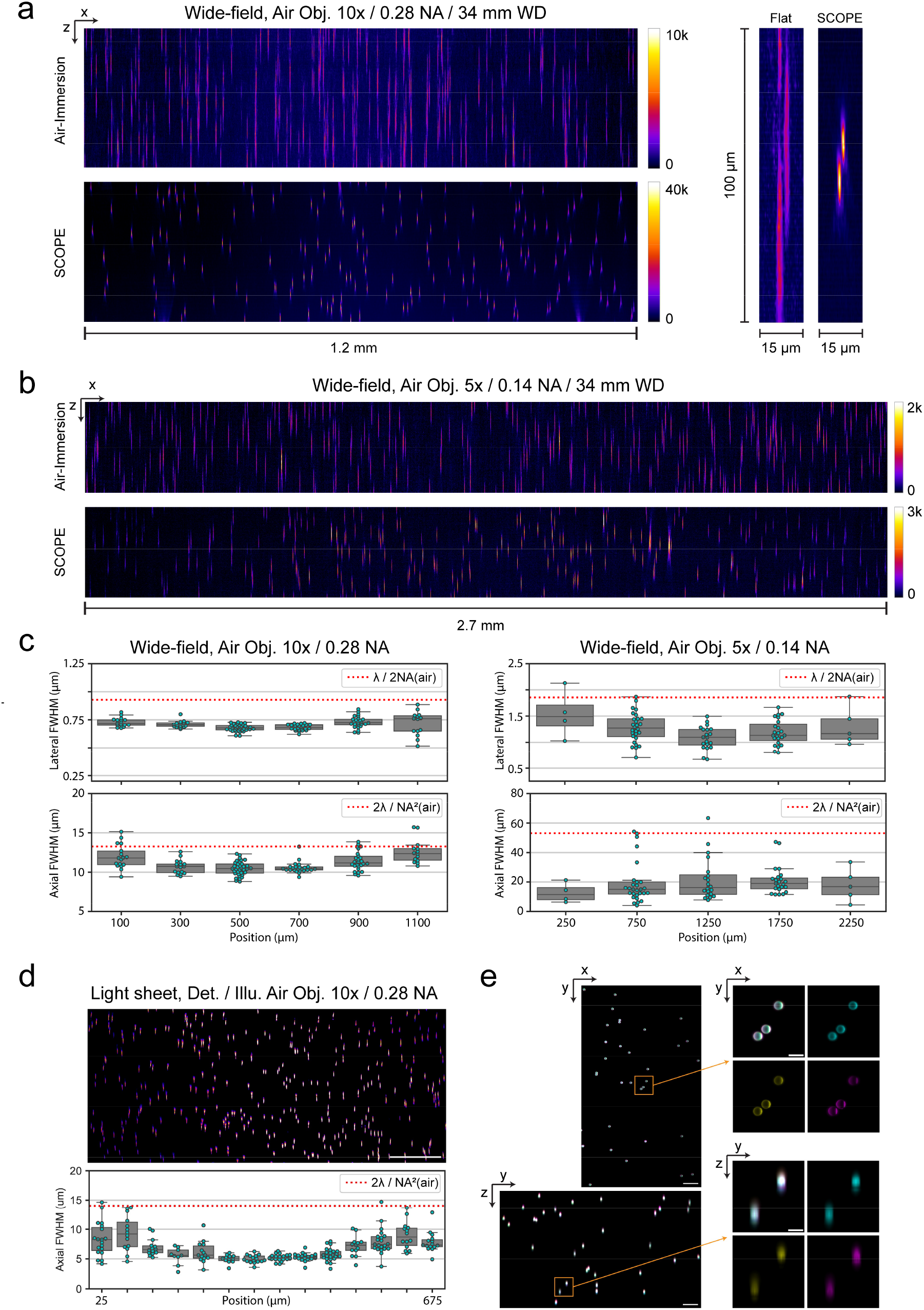
Experimental validation of SCOPE performance through PSF characterization. **(a)** Wide-field imaging of 150 nm gold nanoparticles embedded in 0.5% agarose (H2O) using a 10x/0.28 NA/34 mm WD air objective. Comparison between conventional air-immersion imaging (top) and SCOPE (bottom) configurations shows correction of optical aberrations across a 1.2 mm field of view (FOV) and substantially higher detected signal intensity under identical illumination power. Insets: Representative subregion showing the improved wide-field PSF achieved with SCOPE. **(b)** Equivalent comparison using a 5x/0.14 NA/34 mm WD air objective across a 2.7 mm FOV, demonstrating aberration correction and enhanced signal detection with SCOPE. **(c)** Quantification of lateral and axial full width at half maximum (FWHM) for both objectives across the FOV. SCOPE achieves lateral resolutions of ~0.75 µm (10x) and ~1.2 µm (5x), and axial FWHM values of ~11 µm and ~18 µm, respectively. Dashed red lines indicate theoretical diffraction-limited (Abbe) performance for these air objectives. **(d)** SCOPE-enabled light-sheet microscopy detection PSF characterization using the 10x/0.28 NA/34 mm WD air objective. **(e)** Multicolor light-sheet imaging of polychromatic fluorescent beads demonstrating preserved chromatic alignment across three channels (imaged using 455 nm, 520 nm, and 640 nm illumination wavelengths). Insets show orthogonal (xy, yz, xz) views illustrating co-registered multicolor channels. See also **Supplementary Figure 1b** for a 1 cm-deep three-color imaging stack of polychromatic beads. Scale bars, 100 µm (d), 100 µm (e), and 25 µm (e, insets).

We next quantified the detection PSF by measuring the lateral and axial full width at half maximum (FWHM; **Fig. 2c**), yielding lateral resolutions of ~0.75 µm (10x) and ~1.2 µm (5x), and axial FWHM values of ~11 µm (10x) and ~18 µm (5x). Similar to a classical SIL, the resolution achieved with SCOPE exceeded the theoretical diffraction limits predicted by Abbe’s criterion for the design NA (in air) of these objectives (**Fig. 2c**, red dotted lines) across the entire field of view. Notably, these FWHM values approach those obtained with high-NA immersion objectives, typically orders of magnitude more expensive and limited by much shorter working distances, demonstrating that the hybrid solid-liquid immersion strategy effectively transforms inexpensive air objectives into high-resolution, aberration-free, centimeter-scale imaging systems.

Finally, we integrated the SCOPE imaging device into our low-cost pLSM/SLICE platform and imaged fluorescent beads (**Fig. 2d-e**) to evaluate performance under light-sheet illumination. The resulting data confirmed aberration-free, high-resolution imaging with preserved chromatic alignment across multiple excitation channels (**Fig. 2e**). Moreover, SCOPE maintained high performance for deep imaging, up to 1 cm depth and 2.6 mm field of view shown (**Supplementary Fig. 1d**), demonstrating that its integration into light-sheet microscopy enables standard air objectives to achieve enhanced resolution and achromatic imaging across large volumes while retaining long working distance.

Collectively, these experiments demonstrate that SCOPE enables aberration-free, submicron-resolution imaging across multi-millimeter fields of view and centimeter-scale imaging depths using standard, inexpensive air objectives. Together with low-cost LSM platforms, such as pLSM/SLICE, SCOPE establishes a scalable and cost-effective solution for high-throughput, high-resolution volumetric imaging - a key advance toward democratizing advanced optical microscopy and accelerating AI-driven biological discovery.

### Integration of Expansion Microscopy with SCOPE for Large-Volume, Nanoscale Imaging

Recent advances in ExM^5,6^ enable nanoscale interrogation of biological structures by physically enlarging specimens, achieving effectively sub-diffraction resolution imaging with conventional light microscope optics. However, ExM imposes additional demands: multi-fold expansion dilutes fluorophores, markedly reducing signal per voxel; expanded tissues are mechanically fragile; and intact-sample imaging requires long-working-distance objectives with well-corrected detection and mounting strategies that preserve resolution and signal over large volumes.

Leveraging SCOPE’s ability to pair long-working-distance air objectives with NA enhancement and improved signal collection efficiency, we integrated SCOPE with ExM for nanoscale imaging of large, expanded tissues. Using ~4x expanded *Thy1-GFP* mouse brain sections (see Methods), we systematically optimized final immersion formulation and mounting strategy to balance optical performance, fluorophore stability, and structural integrity while maintaining isotropic expansion. Optimal conditions employed 65-85% glycerol supplemented with ≥2.5 mg/mL DABCO (1,4-diazabicyclo[2.2.2]octane), which preserved fluorophore brightness, minimized photobleaching, and stabilized the expanded tissue. Samples were mounted in quartz cuvettes (10 mm x 10 mm or 10 mm x 20 mm or 20 mm x 20 mm base, scaled as needed) to maintain RI continuity with the SCOPE device.

Under these optimized conditions, expanded *Thy1-GFP* mouse brain sections were imaged using a 10x/0.28 NA/34 mm WD air objective for detection and a 5x/0.14 NA/34 mm WD air objective for light-sheet illumination with the SCOPE-enabled pLSM/SLICE setup. As shown in **Fig. 3** (**Supplementary Video 1**), SCOPE enabled aberration-free, high-resolution imaging of neuronal architecture, clearly resolving dendritic spines and fine neural processes across large tissue volumes. The resulting data were readily compatible with automated reconstruction tools, enabling comprehensive neuronal and dendritic spine tracing (**Fig. 3b**). Given the measured lateral PSF (~0.75 µm; **Fig. 2c**) and ~4x expansion, the effective pre-expansion resolution was ~190 nm, sufficient for automated neurite and spine quantification (**Fig. 3b**).

**Figure 3.**
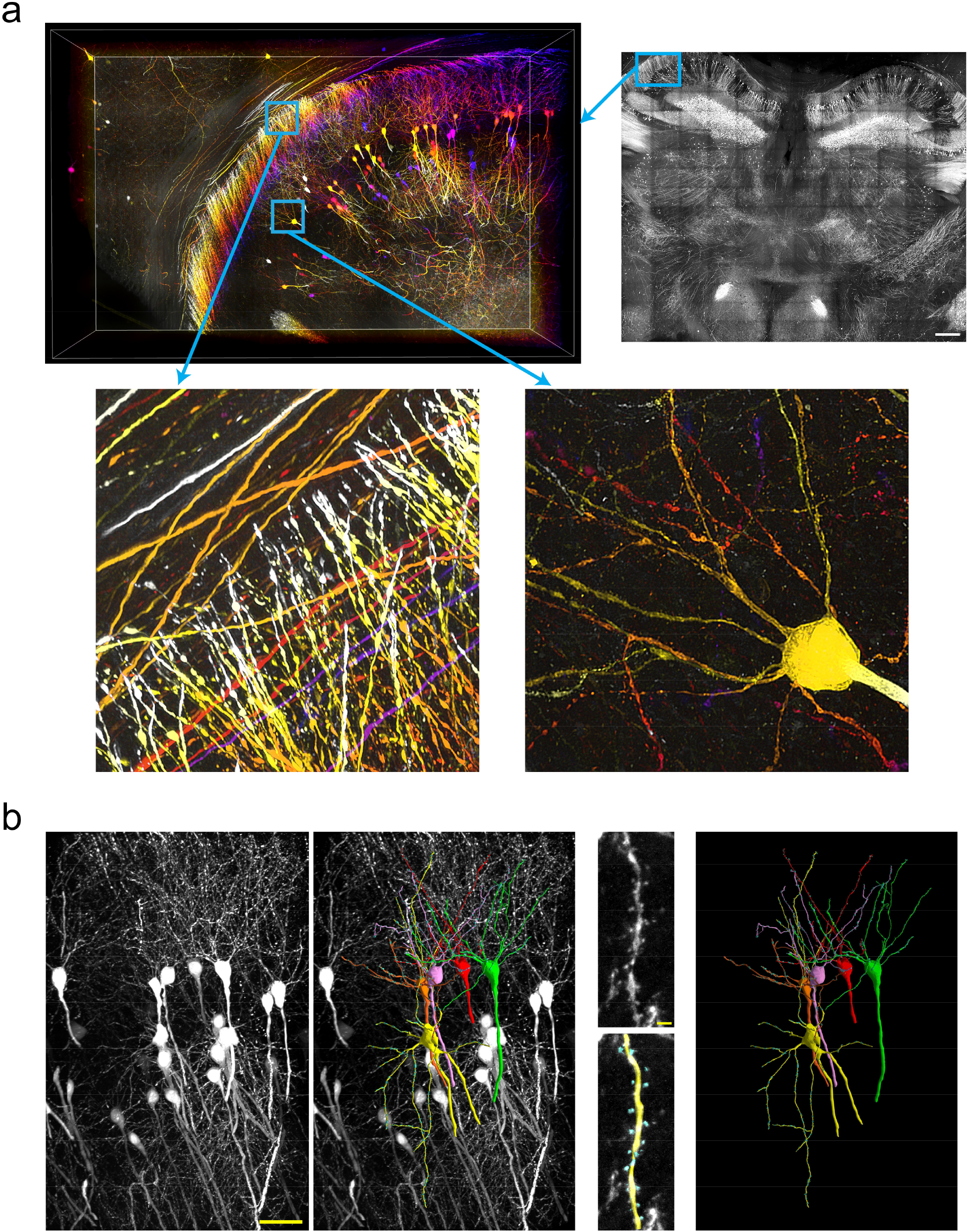
High-resolution volumetric imaging of expanded mouse brain. **(a)** Light-sheet imaging of a ~4x expanded *Thy1-GFP* mouse brain section using the SCOPE imaging device. Left: 3D depth color-coded rendering of a selected region of interest within the hippocampal area, corresponding to the whole-section dataset shown on the right. Insets (bottom): magnified views reveal fine axonal and dendritic processes, including spines. See also **Supplementary Video 1** for a detailed visualization of the dataset. **(b)** Automated tracing of individual neurons, dendritic arbors, and dendritic spines. Left: raw fluorescence image; middle: overlay of data and reconstructed traces; right: isolated traced neuron showing detailed spine morphology. Insets: magnified views of dendritic shafts highlighting individual spines. Scale bars, 1 mm (a, upper-right panel), 100 µm (b, left), and 5 µm (b, insets).

We next developed the ExM-SCOPE workflow for whole intact salamander (*Pleurodeles waltl*) brains, an emerging model in regenerative and comparative neurobiology^20^. By optimizing gelation, digestion, and immersion, we achieved robust 4x isotropic expansion with preserved morphology (Methods). Brains immunostained for tyrosine hydroxylase (TH) were imaged on the SCOPE-enabled pLSM/SLICE, yielding aberration-free, high-resolution volumetric datasets of the entire expanded brain and revealing both mesoscale organization and nanoscale anatomical detail across the entire dopaminergic system (**Fig. 4, Supplementary Videos 2-3**).

**Figure 4.**
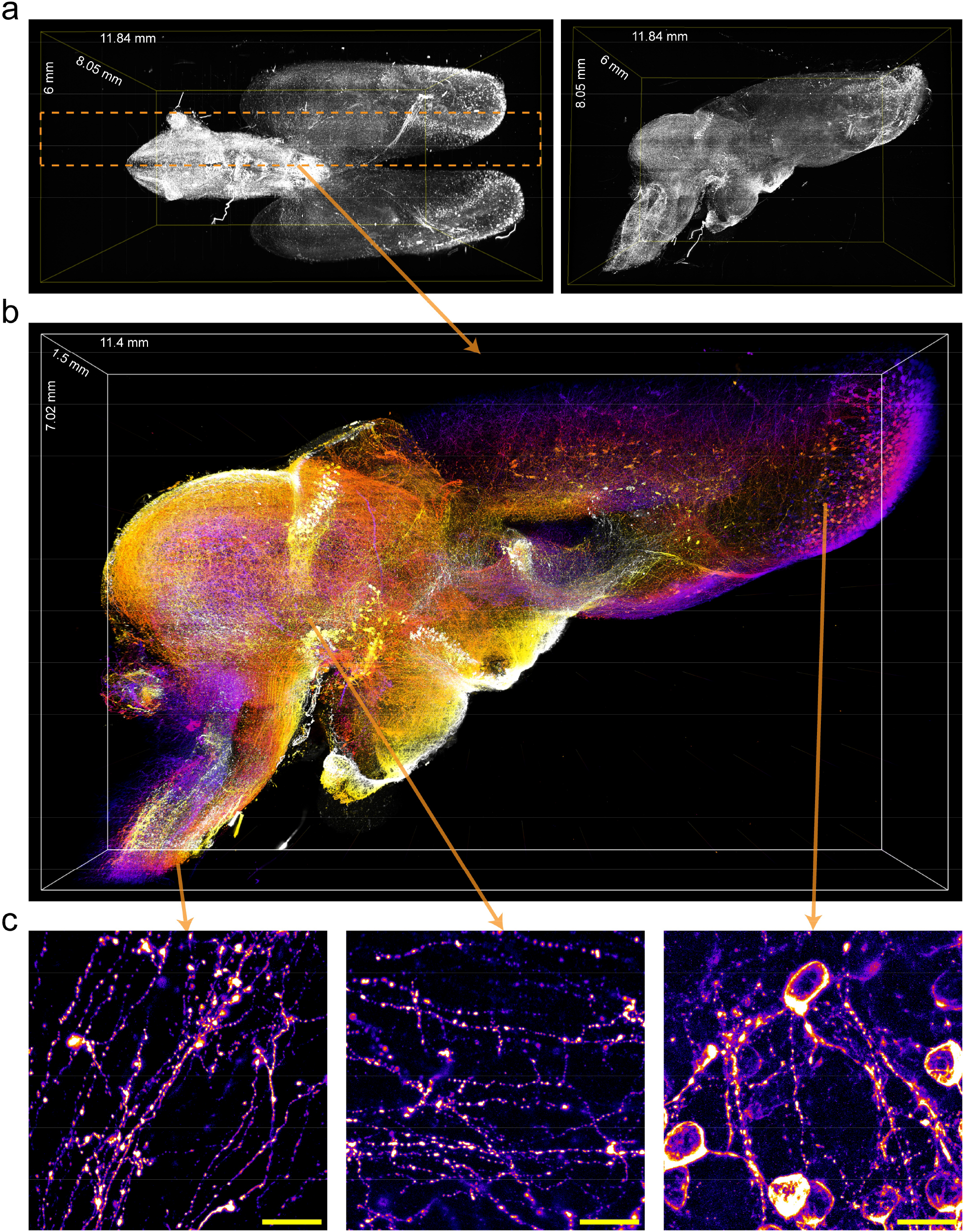
Brain-wide, high-resolution imaging of the dopaminergic system in the salamander brain. **(a)** Volumetric rendering of a whole-brain dataset from an adult salamander (~4x expanded), immunolabeled for tyrosine hydroxylase (TH) and imaged with SCOPE-enabled pLSM/SLICE with 10x/0.28 NA/34 mm WD air objective. Two orthogonal views are shown. See also **Supplementary Videos 2** and **3. (b)** Depth color-coded, high-resolution 3D rendering of a selected region of interest, corresponding to the orange dotted area in (a). **(c)** Maximum-intensity projections of raw data from the hindbrain, midbrain, and forebrain regions, revealing fine neuronal features across the dopaminergic network. Scale bars, 50 µm.

Together, these results demonstrate that SCOPE’s aberration correction, resolution enhancement, and long-working-distance optics with cost-effective light-sheet microscopy platforms such as pLSM enables scalable, high-resolution imaging of large expanded tissues. The combined SCOPE-ExM framework provides a practical route to organ-level nanoscale mapping with low-cost, high-throughput, portable instrumentation.

### SCOPE Enables Multi-Immersion Imaging Across Diverse Clearing Methods

Advanced tissue-clearing methods exhibit wide variability in their optical properties: aqueous-based protocols such as CLARITY^21,22^ yield RIs of ~1.44-1.48, whereas solvent-based methods such as iDISCO^23^ and ECi^24^ reach RI values approaching 1.56. High-quality, aberration-free imaging across this RI range typically requires different specialized objectives or hardware modifications.

Optical simulations of SCOPE (system RI ~ 1.46; **Fig. 1g**) indicated broad tolerance to variations in sample RI while maintaining effective aberration correction across an RI range of 1.38-1.54. This performance eliminates the need for hardware adjustments and supports a single, generalizable workflow for diverse biological and clinical specimens. To experimentally assess this versatility, we evaluated SCOPE (implemented at RI ~ 1.46) across samples immersed in media spanning from aqueous to organic-solvent conditions.

As shown in **Supplementary Fig. 1d**, SCOPE enabled color-corrected deep imaging (up to 1 cm shown) of polychromatic fluorescent beads embedded in 0.5% agarose/H_2_O (RI ~ 1.33), despite the large RI mismatch between the aqueous sample and the hybrid solid-liquid immersion lens implemented with RI ~1.46. We next applied SCOPE to image CUBIC-R–cleared mouse brains (RI ~ 1.51) from the fosTRAP2-tdTomato transgenic line. As shown in **Fig. 5a** and **Supplementary Video 4**, the SCOPE approach provided high-contrast, aberration-free imaging throughout the entire centimeter-scale brain volume (21.13 mm x 16.98 mm x 9.55 mm; x, y, z). The resulting datasets were directly compatible with standard neuroinformatics pipelines, including registration to the Allen Brain Atlas^25^ and automated segmentation and quantification of tdTomato-positive neurons, revealing the expected brain-wide activation patterns. Next, we imaged iDISCO-cleared (RI ~ 1.56) cavefish (*A. mexicanus*) brains stained with nuclear dye TO-PRO-3 (**Fig. 5b** and **Supplementary Video 5**). SCOPE-enabled imaging yielded high-contrast, single-nucleus-resolved images throughout the entire brain, including densely labeled regions (insets, **Fig. 5b**), confirming its effective performance in high-RI, solvent-cleared tissues.

**Figure 5.**
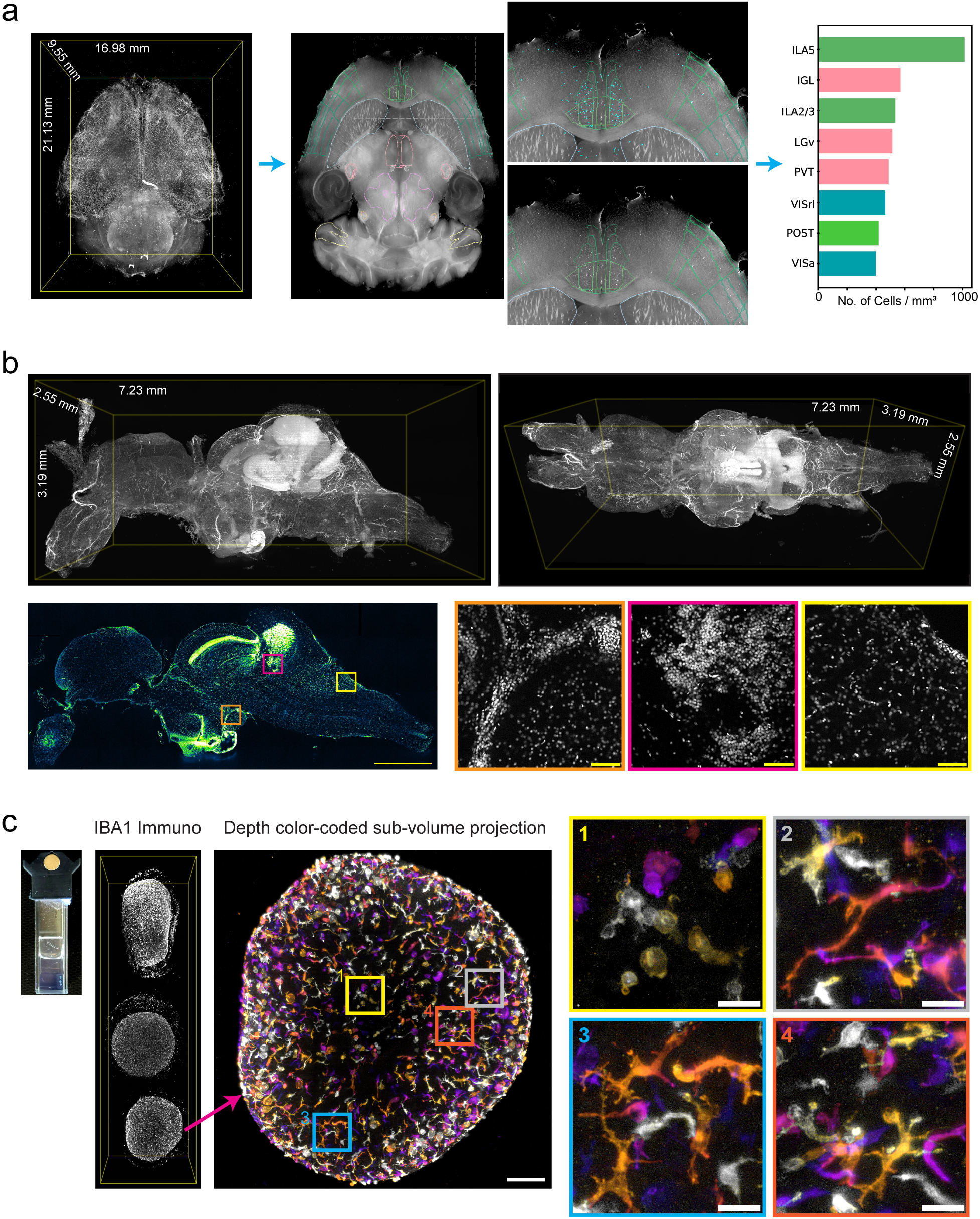
SCOPE enables high-resolution, multi-immersion imaging across diverse tissue clearing methods. **(a)** Whole-brain imaging of a CUBIC-R–cleared *fosTRAP2-tdTomato* mouse brain using the SCOPE-enabled pLSM/SLICE with a 10x/0.28 NA/34 mm WD air objective. Left: Volumetric rendering of the entire mouse brain. Middle: Registration to the Allen Brain Atlas (ABA) and automated segmentation enable anatomical annotation and quantification of labeled neuronal populations. Right: Example showing the top 5% of *tdTomato*+ cells within ABA-defined regions of interest (ROIs). See also **Supplementary Video 4** for the full 3D brain rendering. **(b)** Light-sheet imaging of an iDISCO-cleared, nuclear-labeled (TO-PRO3) *Astyanax mexicanus* (blind cavefish) brain with a SCOPE-enabled 10x/0.28 NA/34 mm WD air objective. Volumetric renderings reveal dense, high-resolution nuclear labeling throughout the brain. Bottom: Sagittal optical section with insets (right) highlighting image quality across distinct, densely labeled brain regions. See also **Supplementary Video 5** for the complete whole-brain dataset. **(c)** High-throughput imaging of human iPSC-derived day-94 cortical organoids co-cultured with microglia and immunolabeled for IBA1, using a SCOPE-enabled 10x/0.28 NA/34 mm WD air objective. Left: Image of a cuvette illustrating parallel imaging of multiple organoids in a single experiment. Center: Depth color-coded sub-volume projection showing extensive microglial coverage and spatial tiling. Right (1-4): Representative magnified views revealing diverse microglial morphologies - amoeboid, elongated, and ramified - consistent with heterogeneous activation states. See also **Supplementary Video 6**. Scale bars, 1 mm (b, bottom left) and 50 µm (b, bottom right).

Further, we demonstrate the high-throughput imaging potential for collections of human iPSC-derived brain organoids with relevance for phenotype screening. SCOPE imaging of Day 94 iPSC-derived cortical organoids incorporated with microglia^26^ (labelled with anti-IBA1) showed complete colonization of the organoid after 2 weeks of co-culture (**Fig. 5c, Supplementary Video 6**). They show wide distribution and tiling throughout the organoid, as well as varying morphology, including amoeboid (**Fig. 5c, sub-panel 1**), elongated (**Fig. 5c, sub-panel 2**), and ramified (**Fig. 5c, sub-panels 3-4**), indicating heterogeneity in inflammatory states.

Together, these results demonstrate that the SCOPE imaging device provides robust, multi-immersion compatibility across tissue-clearing methods ranging from aqueous to solvent-based immersion media without requiring hardware changes, optical realignment, or system reconfiguration.

### Scalable High-Resolution 3D Histopathology with SCOPE

Histopathological analysis remains the cornerstone of preclinical disease research, clinical diagnosis, therapeutic monitoring, and drug discovery. Traditional workflows rely on microscopic examination of thin tissue sections stained with hematoxylin and eosin (H&E) or immunohistochemical markers. While these methods have been indispensable for over a century, they inherently disrupt the native three-dimensional (3D) tissue architecture, limiting insight into spatial cell-cell and cell-matrix relationships that underpin disease progression - a critical limitation in heterogeneous pathologies such as cancer^27^.

Recent studies have demonstrated that deep learning models trained on volumetric histopathological datasets markedly outperform 2D-based approaches for key clinical tasks, including cancer detection, tumor grading, surgical margin assessment, and recurrence prediction^27–29^. These findings underscore the transformative potential of 3D histopathology for precision medicine. Nevertheless, widespread adoption of 3D imaging techniques faces persistent barriers, including prohibitive microscope costs, operational complexity, and limited accessibility. Here, we demonstrate how the SCOPE platform addresses these barriers through its combination of affordability and high-resolution 3D imaging capabilities.

As an initial application, we focused on normal and cancerous human breast tissue. We implemented fluorescent H&E analog labeling of fresh-frozen breast specimens, using TO-PRO-3 for nuclear staining and eosin for cytoplasmic labeling^30^. Samples were cleared and imaged in ECi (RI ~ 1.56), and data were acquired in two channels. Through a false-coloring procedure^31^, the two-channel fluorescence images were converted into conventional H&E-like representations for compatibility with standard pathology workflows.

As shown in **Fig. 6a** and **Supplementary Videos 7-8**, the SCOPE system enables rapid, high-quality 3D imaging of large, intact human breast tissue samples at nuclear resolution. The volumetric datasets reveal the mesoscopic organization of terminal duct-lobular units (TDLUs) into tree-like networks while resolving subcellular structures within individual acini. Imaging of cancerous tissue (**Fig. 6b, Supplementary Video 9**; **Fig. 6c** shows conventional H&E-stained tissue section from the same specimen) further demonstrates the platform’s capability to capture diagnostic histopathological features of malignancy, including abnormal nuclear morphology, fibrotic foci, normal and disrupted tissue architecture, highlighting the potential of SCOPE for scalable, high-resolution 3D histopathology and disease characterization.

**Figure 6.**
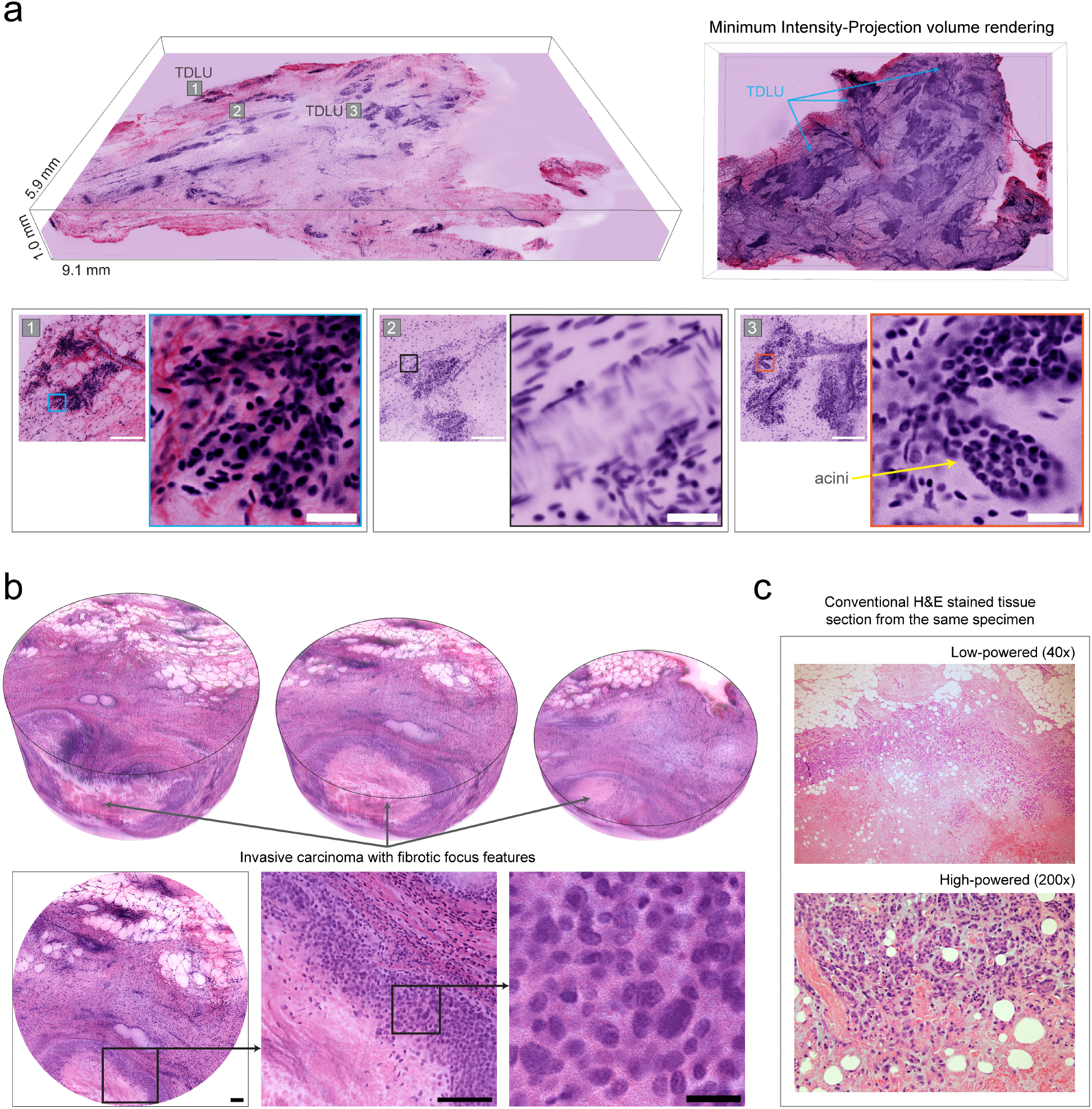
Scalable, high-resolution 3D histopathology of human breast tissue with SCOPE. **(a)** Volumetric rendering of an intact human breast tissue sample imaged using the SCOPE-enabled pLSM/SLICE system with a 10x/0.28 NA/34 mm WD air objective, following fluorescent H&E analog labeling (TO-PRO-3 for nuclei and eosin for cytoplasm), ECi clearing, and false H&E-like false coloring. Renderings with maximum-intensity projection (right) and minimum-intensity projection (left), revealing nuclear-resolution architecture of terminal duct-lobular units (TDLUs) and their mesoscopic organization into tree-like structures. Insets (1-3) show magnified views of regions marked in the maximum-intensity projection rendering. Scale bars, 200 µm (overview) and 25 µm (magnified views). See also **Supplementary Videos 7** and **8** for detailed visualization of the dataset. **(b)** 3D rendering of a cancerous human breast tissue sample showing invasive carcinoma with fibrotic focus features, visualized at three different depth levels. Bottom row: Single optical section with corresponding magnified views as indicated. See also **Supplementary Video 9** for detailed visualization of the dataset. Scale bars (left to right): 100 µm, 100 µm, and 20 µm. (c) Conventional 2D H&E-stained tissue section from the same specimen shown in b, illustrating invasive carcinoma with fibrotic-focus morphology at low and high power.

## DISCUSSION

We present the HySIL optical framework, which enables sophisticated wavefront control through seamless integration of a solid optical element with a precisely RI-matched liquid medium. This hybrid configuration demonstrates exceptional performance when combined with long-working-distance, low-cost air objectives positioned at the solid interface for signal collection, while specimen is positioned in the liquid compartment (housed in RI-matched glass cuvettes) to enable aberration-corrected, high-resolution 3D imaging of large samples. This principle was realized as modular SCOPE and Super-SCOPE imaging devices (**Fig. 1**) that integrate readily into existing open-source and commercial LSM platforms, including the low-cost pLSM^11^-based platforms, yielding a highly compact, portable system for subcellular-resolution imaging across large volumes. Importantly, although our initial implementation of SCOPE employs readily available plano-convex lenses for simplicity and cost-effectiveness, the HySIL framework is broadly generalizable, pairing any solid optical element with an RI-matched liquid, to enable customization of lens geometries and materials for optimized performance across diverse imaging applications.

A key advantage of SCOPE is its remarkable robustness to sample RI variations, maintaining excellent performance across a broad RI range (1.33-1.56) without hardware modifications while preserving chromatic correction and providing wide detection field-of-view (up to 2.6 mm, **Fig. 2b**). We validated this multi-immersion versatility through aberration-free imaging of samples spanning aqueous to organic immersion media, including 1-cm-thick volumes of polychromatic fluorescent beads in 0.5% agarose/H_2_O (**Supplementary Fig. 1b**), ExM-expanded mouse (**Fig. 3**) and salamander (**Fig. 4**) brains, CUBIC-cleared intact mouse brains with centimeter-scale dimensions in all three axes (**Fig. 5a**), iDISCO-cleared cavefish brains (**Fig. 5b**), and high-throughput screening of brain organoids revealing subcellular details of microglial morphology across different activation states (**Fig. 5c**).

Beyond basic scientific research applications, we demonstrated SCOPE as a scalable, benchtop solution for 3D histopathological imaging of clinically relevant specimens (**Fig. 6, Supplementary Videos 7–9**). This capability directly addresses a fundamental limitation of conventional tissue sectioning approaches, which destroy the 3D spatial relationships essential for disease characterization^27^. Our proof-of-concept studies on normal and cancerous human breast tissues achieved nuclear-resolution imaging of large, intact volumes, generating datasets amenable to false-coloring for conventional H&E-like visualizations compatible with existing diagnostic workforces and workflows. The ability to simultaneously resolve mesoscale tissue organization (e.g., the tree-like structure of lobules in breast tissue) and cellular/nuclear features, while remaining affordable and portable, can accelerate advances in computational pathology, machine learning-based diagnostics, and precision medicine.

Despite the robust performance of HySIL and SCOPE, several areas present opportunities for future enhancements. First, although HySIL amplifies the effective NA in proportion to the RI of the hybrid lens, fundamental limits on resolution and light collection are still determined by the NA of the air objective, which generally decreases as the working distance increases. The Super-SCOPE configuration, which employs a Weierstrass super hemispherical geometry^17^, provides a further increase in effective NA that scales with *n*^2^, offering higher resolution but at the cost of tighter optical tolerances and a narrower detection spectral bandwidth to minimize chromatic aberrations. Future work could refine these geometric designs to reduce chromatic errors or incorporate computational post-processing approaches for color correction. In addition, using solid components with even higher refractive indices paired with matching immersion media, in combination with higher NA air objectives (for example, the Mitutoyo Plan Apochromat 20×/0.42 NA/20 mm WD) with slightly shorter working distances, could ultimately achieve or surpass the resolution and photon collection efficiency of the high NA immersion objectives, which typically provide only a few hundred microns of working distance. Second, although we demonstrate compatibility across the most widely used tissue clearing methods, samples with extreme RI values (<1.33 or >1.56) may require alternative HySIL configurations using different optical materials or liquid formulations to maintain aberration correction. Importantly, SCOPE’s modular and easy-to-swap architecture facilitates straightforward use of multiple SCOPE chambers to accommodate extreme RI requirements. Third, while our current implementation is demonstrated for horizontal benchtop LSM systems, the underlying HySIL principles are configuration-independent and can be adapted to upright or inverted configurations through appropriate chamber design.

In summary, the HySIL framework and SCOPE and Super-SCOPE implementations establish a practical and cost-effective foundation for next-generation volumetric microscopy, bridging the gap between cutting-edge optical performance, scalability, and widespread accessibility. This advance marks a significant step toward the democratization of high-resolution biological imaging and the acceleration of data-driven scientific discovery in preclinical research as well as in potential clinical applications in 3D histopathology.

## METHODS

### Design, simulations and PSF characterization of SCOPE

#### Optical simulations

All optical simulations were performed in Ansys Zemax OpticStudio (2024). As illustrated by the ray traces in **Fig. 1e** and **Supplementary Fig. 1a**, the optical model consisted of a paraxial detection lens (serving as the objective), a Thorlabs TTL200 tube lens (black-box model provided by Thorlabs), and a plano-convex fused-silica lens (radius of curvature = 16 mm in **Fig. 1e** or 27.5 mm in **Supplementary Fig. 1a**; Corning 7980 substrate) serving as the solid element of the hybrid immersion lens. The RI of the immersion liquid was set to match that of the fused silica (n ~ 1.46), forming a continuous solid-liquid optical system. The sample was modeled as a planar slab of defined thickness and RI.

To enable direct comparison across configurations, air-air, air-immersion, SCOPE, and Super-SCOPE, a fixed aperture stop was used to represent the same detection objective in all models. The aperture diameter was set according to an air objective with NA = 0.28, matching the Mitutoyo 10×/0.28 NA objective used experimentally. For the air-immersion configuration, the distance between the flat glass window and the object plane being imaged was adjusted based on the optical path difference in oil to ensure equivalent optical and geometrical distances. In the SCOPE configuration, the distance between the convex surface of the correction lens and the object plane being imaged was set equal to the lens radius (r). For Super-SCOPE, this distance followed r (1 + 1/*n*), where r is the lens radius and *n* is the refractive index of the lens material. The fast Semi-Diameters option in Zemax was used to ensure that the aperture was calculated using marginal rays for high accuracy. The field type was defined as an object height corresponding to a 700 µm field of view (FOV) at the sample plane. Simulations were performed for three representative wavelengths, 520 nm, 590 nm, and 640 nm, with relative weights of 5.0, 1.0, and 1.0, respectively. The objective position was optimized as a variable within +/−2 mm of its nominal position in all configurations.

The Huygens PSF, its cross-section, and the corresponding diffraction-based encircled-energy plots (shown in **Fig. 1e–f**) were calculated at 520 nm for each configuration, with pupil and image sampling set to 512 x 512. The relative irradiance, normalized to the Strehl ratio, was calculated, and detection-plane distances were scaled by the system magnification: 5x (air), 4.9x (air-immersion), 7.3x (SCOPE), and 10.7x (Super-SCOPE). The optical path difference (OPD) wavefront error, used to quantify residual aberrations (**Supplementary Fig. 1b**), was evaluated at 520 nm with NA = 0.28 (corresponding to the Mitutoyo 10x objective) over a 700 µm FOV. Tangential and sagittal planes were analyzed along orthogonal field axes to capture off-axis aberrations, and results were compared between the center and edge of the FOV for SCOPE and conventional flat-window (air-immersion) configurations. The root-mean-square (RMS) wavefront error (**Fig. 1g, Supplementary Fig. 1c**) was computed at 520 nm while varying both sample RI and sample thickness. The system was re-optimized for each condition to maintain optimal performance. RI values were varied in increments of +/−0.02 around the system baseline (n = 1.46).

#### SCOPE device Implementation

The SCOPE imaging device consisted of an off-the-shelf plano-convex lens (Edmund Optics cat #48-670; R = 27.5 mm, Ø = 25 mm) as the solid component and a Cargille Laboratories RI-matching immersion oil (n ~ 1.46, Code: 50350) as the liquid component, forming a hybrid solid-liquid seamless system. The chamber body was 3D-printed using black resin and sealed with optical-grade epoxy to ensure mechanical stability and prevent leakage. The lens was fixed to the chamber window using UV-curable adhesive. For detection, Mitutoyo 10x/0.28 NA and 5x/0.14 NA long WD air objectives were used; illumination was provided by Mitutoyo 5x/0.14 NA for most experiments.

#### PSF Imaging and Characterization

To measure the system PSF, gold nanoparticles (Sigma-Aldrich, #742058; 150 nm diameter) were suspended in 0.5% agarose to a final concentration of 5% (v/v) in 1.5 mL total volume. For wide-field PSF imaging, the pLSM illumination light sheet was expanded to full extent to result in illumination of entire volume. The scattered light was collected in the detection arm without emission filter. Z-stacks were acquired at 2 µm step size using identical laser power for all conditions to enable quantitative signal comparison. For fluorescence-based PSF measurements and analysis, 15 µm polychromatic (FocalCheck (TM) Microspheres, Thermo Fisher Scientific, cat. number: F7239) and 1 µm fluorescent beads (Polysciences, Fluoresbrite Microparticles, cat. number: 18660) were embedded in 0.5% agarose under identical conditions. Appropriate excitation and emission filters were selected for each spectral channel.

#### PSF FWHM Analysis

Image stacks were analyzed to determine the lateral (X-Y) and axial (Z) FWHM of individual PSFs. All beads or nanoparticles in an image volume were manually marked by using a custom Python-based GUI from maximum-intensity projection image. Center coordinates were further refined by locating the pixel of maximum intensity in the immediate neighborhood. Intensity profiles along the X, Y, and Z axes were extracted and fit to a Gaussian model, from which FWHM values were calculated. The resulting lateral and axial resolutions were compared across objectives and configurations (SCOPE vs. Air-Immersion).

### Sample Preparations

#### Expansion microscopy of Salamander brain

Building upon multiple published expansion microscopy (ExM) protocols, we systematically optimized experimental parameters to establish a reliable method for whole-brain expansion in the salamander *Pleurodeles waltl*, as described below.

An adult salamander was obtained from a breeding colony established at Columbia University, with all experiments conducted in accordance with the National Institute of Health (NIH) guidelines and Columbia University institutional animal care and use committee policies governing animal use and welfare. The animal was anesthetized by immersion in 0.1% MS-222, followed by transcardial perfusion with 10 mL of cold 1x phosphate-buffered saline (PBS) to flush out the blood, and then with 10 mL of cold 4% paraformaldehyde (PFA) in PBS for fixation. The brain was carefully extracted and post-fixed overnight in 4% PFA at 4°C. Whole-brain immunolabeling was performed according to the salamander immunohistochemistry protocol described previously^32^. To label dopaminergic neurons and their processes, we used a rabbit anti-tyrosine hydroxylase primary antibody (1:1000, Sigma-Aldrich AB152), followed by a goat anti-rabbit Alexa Fluor 546-conjugated secondary antibody (1:1000, Fisher A-11035).

Following immunostaining, the sample was transferred from PBS to MES-buffered saline [2-(N-morpholino)ethanesulfonic acid] to enhance penetration of the anchoring reagent. This step was adapted from a whole-brain ExM protocol for mouse brain^33^. The brain was incubated in Acryloyl-X (AcX) at 4°C for 48 hours - adjusted for the smaller size of the salamander brain. The sample was then washed twice in PBS at 4°C for 1 hour each, followed by an overnight wash. For hydrogel embedding, the Stock X monomer solution was adapted from the TissUExM protocol^34^, with increased acrylamide concentration based on our optimization experiments. The final monomer composition was: 23% sodium acrylate, 10% acrylamide, 0.1% N,N′-methylenebisacrylamide, and 1x PBS. This solution was activated with 0.12% (w/v) VA-044, and the brain was incubated in the activated monomer mix at 4°C for 2 days.

For gelation, the sample was degassed in a vacuum chamber for 20 minutes, purged with nitrogen gas, and then incubated in an airtight container at 37°C for approximately 3 hours, or until complete polymerization was observed^35^. The gelled brain was then removed from the container, trimmed, and briefly washed twice in PBS (5 minutes each). The embedded sample was incubated in digestion buffer containing 8 U/mL of Proteinase K at room temperature for 24 hours, or until the tissue appeared fully cleared. Following digestion, the gel was washed twice in PBS (1 hour each), followed by an overnight wash at room temperature.

Expansion was initiated by immersing the sample in deionized water, with water changes every 30-60 minutes. Expansion progress was monitored visually and halted once the desired expansion factor was achieved. The sample was then re-immersed in PBS to stabilize the gel and reduce over-expansion. To better match the RI of the expanded sample for optimal imaging, the tissue was equilibrated in a graded series of glycerol/PBS solutions, culminating in 65% glycerol in PBS (RI ~ 1.42). To prevent photobleaching during imaging and long-term storage, 0.5% DABCO (1,4-diazabicyclo[2.2.2]octane) was added to all glycerol solutions. The fully expanded, RI-matched sample was imaged using the pLSM/SLICE microscope equipped with the SCOPE imaging device.

#### Expansion microscopy of mouse brain sections

These mouse procedures were approved by the Columbia University Institutional Animal Care and Use Committee (IACUC, protocol #AC-AABG0556). An adult (8-10 weeks old) transgenic *Thy1-GFP* (C57BL/6) mouse was transcardially perfused with 4% PFA, and the brain was extracted and post-fixed overnight in 4% PFA at 4°C. The brain was then washed in PBS three times for 30 minutes each. A 400 μm-thick vibratome section was prepared for expansion microscopy. For processing, the same overall protocol described for the salamander brain was used, with adjustments to incubation times to account for the reduced tissue thickness. Specifically, AcX treatment was carried out overnight in PBS instead of MES-buffered saline, as the lower pH was not required for thin section penetration^33^. The brain section was then incubated in activated Stock X monomer solution for 1 hour at 4°C, followed by gelation and digestion as described above. Digestion was carried out for 24 hours at room temperature in the dark. The embedded sample was expanded by immersion in deionized water for a total of 3 hours, with water changes every 30-60 minutes and was subsequently returned to PBS. For imaging, the sample was RI-matched using the same 65% glycerol in PBS solution with 0.5% DABCO and imaged using the pLSM/SLICE microscope with SCOPE imaging device.

#### CUBIC-R clearing of FosTRAP whole mouse brain

FosTRAP transgenic mice^36^, aged 3-5 months, were exposed to a novel complex environment for 1 hour to induce neuronal activity. Immediately following this exposure, mice received an intraperitoneal injection of 4-hydroxytamoxifen (4-OHT; 50 mg/kg). One week later, animals were transcardially perfused with saline followed by 4% formaldehyde. Brains were extracted, post-fixed in 4% formaldehyde for 24 hours at 4°C, and then transferred to PBS. Tissue clearing was performed using a modified version of the CUBIC-R protocol^37^. Neurons that expressed *Fos* during the 1-hour stimulation period prior to tamoxifen administration were permanently labeled with tdTomato, enabling visualization of activity-tagged neuronal populations across the entire brain. Cleared brains were imaged using the SCOPE-enabled pLSM/SLICE microscope. All animal procedures were conducted in accordance with an IACUC-approved protocol from the National Institute of Mental Health (NIMH; protocol #LSN-02).

#### iDISCO cleared cavefish brains

Adult *Astyanax mexicanus* were euthanized and decapitated at the gill plate using a scalpel, and the heads were briefly immersed in ice-cold 1x phosphate-buffered saline (PBS) for 30 seconds to remove residual blood. Samples were then fixed in 4% paraformaldehyde (PFA) in 1x PBS at 4°C overnight (18-20 hours) with gentle shaking. The following day, heads were washed three times in 1x PBS at room temperature (RT) for 30 minutes each, with shaking. Brains were dissected and dehydrated through a graded methanol/Milli-Q water series (20%, 40%, 60%, 80%, 100%, 100%) and stored in 100% methanol at 4°C until further processing for iDISCO+ staining, as in^23,38,39^.

For staining, brains were bleached overnight at 4°C in 5% hydrogen peroxide in methanol without shaking (16-18 hours). The next day, samples were rehydrated through a reverse methanol/Milli-Q water gradient (80%, 60%, 40%, 20%) and finally placed into 1x PBS. Samples were then washed twice in 0.2% Triton X-100 in PBS (PBSTx) for 30 minutes each at RT with rocking. Brains were permeabilized in a solution of 2.3% glycine and 20% dimethyl sulfoxide (DMSO) in PBSTx at 37°C for 24 hours without shaking. Blocking was performed in 6% normal donkey serum and 10% DMSO in PBSTx at 37°C for an additional 24 hours without shaking. Samples were then incubated in TO-PRO-3 iodide nuclear stain (Invitrogen, Cat#: T3605) at a 1:10,000 dilution in PTwH buffer (0.1% heparin, 0.2% Tween-20 in 1x PBS) containing 3% normal donkey serum and 5% DMSO, for 24 hours at 37°C without shaking. After staining, brains were washed 4-5 times in PTwH over the course of a day at RT on a rocker. This wash step was repeated the following day under the same conditions, with samples protected from light, as in^23,38,39^.

For clearing, the brains were dehydrated again through a methanol/Milli-Q water gradient (20%, 40%, 60%, 80%, 100%, 100%) for 30 minutes per step at RT on a rocker. Samples were then incubated in 66% dichloromethane (DCM) in methanol for 3 hours at RT with rocking, followed by two 15-minute washes in 100% DCM. Final clearing was performed in dibenzyl ether (DBE) for at least 24 hours at RT in the dark. All tubes were filled completely with DBE to prevent air exposure during storage and imaging. The cleared sample was mounted in a cuvette and imaged with pLSM/SLICE equipped with SCOPE imaging device.

#### Preparation of iPSC-derived cortical organoids with microglia

Cortical organoids with microglia were generated from an hiPSC line derived from a healthy male under Columbia University IRB protocol AAAP0052 (Lopes da Costa *et. al*)^40^. On Day 0, 9,000 iPS cells were seeded per well in a 96-well U-bottom plate in DMEM/F12, 15% KSR, 5% FBS, 1% NEAA, 1% Glutamax, 100uM beta-mercaptomethanol, 100nM LDN-193189, 10uM SB-431542, 2uM XAV-939, and 50uM Y27632. On day 2, the media was replenished without Y27632. On days 4-8, the media was replenished every other day without FBS. On day 10, the media was changed to 50% DMEM/F12, 50% Neurobasal, 0.5% N2, 1% B27 without RA, 0.5% NEAA, 1% Glutamax, 0.025% insulin, 1% pen-strep, and 50uM beta-mercaptoethanol, replenished every other day through day 16. On day 18, the media was changed to 50% DMEM/F12, 50% Neurobasal, 0.5% N2, 1% B27 with vitamin A, 0.5% NEAA, 1% glutamax, 0.025% insulin, 50uM beta-mercaptoethanol, 1% pen-strep, 20ng/mL BDNF, and 200uM ascorbic acid. The media was replenished every other day until day 26, after which the media was changed every 4-5 days. Microglial progenitors were generated as previously described^26^. Progenitors were harvested and co-cultured with day 80 cortical organoids at 50,000 cells/organoid in IL-34 (100ng/mL) and M-CSF (10 ng/mL) for 2 weeks before analysis.

Organoids were fixed in 4% PFA overnight and washed with PBS 3x for 10 minutes, followed by simultaneous blocking and permeabilization in 5% serum/0.5% TritonX in PBS for 2 hr at room temperature. The organoids were incubated with chicken anti-IBA1 (Synaptic Systems) at 1:500 in 5% serum/0.1% Triton-X overnight, were washed in 0.05% Tween-20/PBS 3x for 10 minutes and incubated with goat anti-chicken 647 secondary antibody in 5% serum/0.1% Triton-X overnight. Finaly, the organoids were washed 0.05% Tween-20/PBS 3x for 10 minutes and stored in PBS with azide.

Organoids were aligned in 1% agarose and cleared using F-disco as previously described^11,41^. They were placed sequentially for 1 hour at 4^°^C in 50% THF/dH20, 70% THF/dH20, 80% THF/dH20, 100% THF/dH20, 100% THF/dH20, and 100% dibenzyl ether. Cleared organoids were stored in 100% dibenzyl ether at 4^°^C.

#### 3D Histopathology: Fluorescent H&E-Like Staining and Clearing of Human Breast Tissue

Frozen, de-identified human breast tissue specimens were obtained from the Tumor Bank (IRB AAAB2667) at the Herbert Irving Comprehensive Cancer Center, Columbia University, and stored at −80 °C. Samples were thawed at room temperature (RT) for 30 minutes and pre-washed in 70% ethanol (v/v in deionized water) to remove residual cryoprotectant, and immersed in 70% ethanol within 50 mL Falcon tubes, with shaking on an orbital shaker at 60-80 rpm for a minimum of 3 hours at RT.

For fluorescent H&E-like staining^30^, a staining buffer was prepared consisting of 70% ethanol, 10 mM NaCl, and adjusted to pH 4 using 0.1 N HCl. TO-PRO-3 (1:500 v/v) and eosin (1:100 v/v) were diluted into this buffer to prepare the staining solution. Samples were incubated in the staining solution for 48 hours at RT with gentle agitation. Following staining, tissues were dehydrated by incubation in 100% ethanol for 1 hour at RT, followed by a second incubation in fresh ethanol overnight to ensure complete dehydration. For optical clearing, samples were immersed in 10 mL of ethyl cinnamate (ECi) for 2 hours at RT with gentle agitation, followed by replacement with fresh ECi and an additional 2-hour incubation. For imaging, samples were transferred into 10 x 10 mm cuvettes containing 1.5 mL of ECi and imaged using the pLSM/SLICE microscope equipped with the SCOPE imaging device.

### Imaging experimentations

All imaging experiments were performed using the SCOPE imaging device which incorporates a hybrid solid-liquid immersion lens system composed of a fused quartz plano-convex lens with a 27.5 mm radius of curvature and immersion oil matched to a RI of ~1.46. SCOPE was integrated into the lab-built pLSM and SLICE system (MBF Bioscience), a commercial implementation of the pLSM design. Samples were mounted in quartz cuvettes (10 x 20 mm or 10 x 10 mm base dimensions) to ensure RI continuity across optical interfaces. Image detection was carried out using Mitutoyo long WD air objectives (10x/0.28 NA/34 mm WD; 5x/0.14 NA/34 mm WD). For most experiments, a 5x/0.14 NA/34 mm WD illumination objective was used. Multicolor imaging was performed using laser excitation at wavelengths of 455 nm, 520 nm, and 640 nm. Appropriate emission bandpass filters were selected based on the spectral characteristics of the fluorophores used.

### Data analysis and visualization

All stitching and visualization of imaging data were performed using BrightSLICE software (MBF Bioscience). Detailed volumetric renderings were generated using ImageJ/FIJI^42–44^, Neurolucida 360, and Amira (Thermo Fisher Scientific). Automated neuronal tracing and 3D reconstructions were carried out with Neurolucida 360. Whole-brain registration and cellular quantification were performed using NeuroInfo software (MBF Bioscience).

## ACKNOWLEDGEMENTS

We thank the members of the Tomer Lab for their discussions and input. We are grateful to the families who donated tissues to the Tumor Bank at the Herbert Irving Comprehensive Cancer Center, Columbia University, for use in this study. We thank Sage Aronson for early discussions related to the SCOPE imaging device. R.T. acknowledges partial support for the research described in this study from the NIH (grant number DP2MH119423) and a Columbia University Arts and Sciences startup grant. S.R.G. was supported by the Leon Levy Scholarship in Neuroscience and the NIH (grant number R25MH086466). J.G. acknowledges partial support for the research described in this study from the NIH (award number 42MH124566). R.M. and J.E.K acknowledge support from the NIH (grant number R35GM138345). C.R.G. acknowledges support from the NIMH-IRP (ZIA MH002497-36). R.H. acknowledges support from Hope for Depression Research Foundation (HDRF) grant RGA13. M.A.T. acknowledges support from the NIH (grant number 1U01MH139723). This research was funded in part through the NIH/NCI Cancer Center Support Grant P30CA013696 and used the Molecular Pathology Shared Resource (MPSR) Tissue Bank. The content is solely the responsibility of the authors and does not necessarily represent the official views of the National Institutes of Health.

## AUTHOR CONTRIBUITONS

R.T. conceived and designed the HySIL/SCOPE/Super-SCOPE. M.F., E.N., B.H., C.G., J.G. and R.T. implemented the SCOPE imaging device and integrated into pLSM/SLICE systems. C.G. and R.T. developed the Zemax optical analysis, performed beads imaging and contributed to the imaging of biological samples. Development, preparation, and imaging of biological samples were performed as follows: P.A., E.D.D., and R.T. for mouse and salamander ExM, with salamander resources and expert inputs from M.A.T.; C.R.G. and M.F. for CUBIC-R mouse brain; R.M., M.F., and J.K. for cavefish brain; S.R.G., R.H., and C.D.M. for brain organoids with microglia; and C.G., E.D.D., H.H., and R.T. for 3D histopathology. R.T., N.J.O., M.H., M.F., and J.G. performed the data analysis. R.T. wrote the manuscript with input from all authors and performed overall supervision of the project.

## REFERENCES

1. Jumper, J., Evans, R., Pritzel, A., Green, T., Figurnov, M., Ronneberger, O., Tunyasuvunakool, K., Bates, R., Žídek, A., Potapenko, A., et al. (2021). Highly accurate protein structure prediction with AlphaFold. Nature 596, 583–589. 10.1038/s41586-021-03819-2.

2. Hayes, T., Rao, R., Akin, H., Sofroniew, N.J., Oktay, D., Lin, Z., Verkuil, R., Tran, V.Q., Deaton, J., Wiggert, M., et al. (2025). Simulating 500 million years of evolution with a language model. Science 387, 850–858. 10.1126/science.ads0018.

3. van der Laak, J., Litjens, G., and Ciompi, F. (2021). Deep learning in histopathology: the path to the clinic. Nat Med 27, 775–784. 10.1038/s41591-021-01343-4.

4. Bunne, C., Roohani, Y., Rosen, Y., Gupta, A., Zhang, X., Roed, M., Alexandrov, T., AlQuraishi, M., Brennan, P., Burkhardt, D.B., et al. (2024). How to build the virtual cell with artificial intelligence: Priorities and opportunities. Cell 187, 7045–7063. 10.1016/j.cell.2024.11.015.

5. Chen, F., Tillberg, P.W., and Boyden, E.S. (2015). Expansion microscopy. Science 347, 543–548. 10.1126/science.1260088.

6. Wassie, A.T., Zhao, Y., and Boyden, E.S. (2019). Expansion microscopy: principles and uses in biological research. Nat Methods 16, 33–41. 10.1038/s41592-018-0219-4.

7. Stelzer, E.H.K., Strobl, F., Chang, B.-J., Preusser, F., Preibisch, S., McDole, K., and Fiolka, R. (2021). Light sheet fluorescence microscopy. Nat Rev Methods Primers 1, 1–25. 10.1038/s43586-021-00069-4.

8. Pitrone, P.G., Schindelin, J., Stuyvenberg, L., Preibisch, S., Weber, M., Eliceiri, K.W., Huisken, J., and Tomancak, P. (2013). OpenSPIM: an open-access light-sheet microscopy platform. Nat Methods 10, 598–599. 10.1038/nmeth.2507.

9. Voigt, F.F., Kirschenbaum, D., Platonova, E., Pagès, S., Campbell, R.A.A., Kastli, R., Schaettin, M., Egolf, L., van der Bourg, A., Bethge, P., et al. (2019). The mesoSPIM initiative: open-source light-sheet microscopes for imaging cleared tissue. Nat Methods 16, 1105–1108. 10.1038/s41592-019-0554-0.

10. Vladimirov, N., Voigt, F.F., Naert, T., Araujo, G.R., Cai, R., Reuss, A.M., Zhao, S., Schmid, P., Hildebrand, S., Schaettin, M., et al. (2024). Benchtop mesoSPIM: a next-generation open-source light-sheet microscope for cleared samples. Nat Commun 15, 2679. 10.1038/s41467-024-46770-2.

11. Chen, Y., Chauhan, S., Gong, C., Dayton, H., Xu, C., De La Cruz, E.D., Tsai, Y.-Y.W., Datta, M.S., Rosoklija, G.B., Dwork, A.J., et al. (2024). Low-cost and scalable projected light-sheet microscopy for the high-resolution imaging of cleared tissue and living samples. Nat. Biomed. Eng 8, 1109–1123. 10.1038/s41551-024-01249-9.

12. Turcotte, R., Liang, Y., and Ji, N. (2017). Adaptive optical versus spherical aberration corrections for in vivo brain imaging. Biomed. Opt. Express, BOE 8, 3891–3902. 10.1364/BOE.8.003891.

13. Barner, L.A., Glaser, A.K., True, L.D., Reder, N.P., and Liu, J.T.C. (2019). Solid immersion meniscus lens (SIMlens) for open-top light-sheet microscopy. Opt. Lett., OL 44, 4451–4454. 10.1364/OL.44.004451.

14. Glaser, A., Chandrashekar, J., Vasquez, S., Arshadi, C., Javeri, R., Ouellette, N., Jiang, X., Baka, J., Kovacs, G., Woodard, M., et al. (2025). Expansion-assisted selective plane illumination microscopy for nanoscale imaging of centimeter-scale tissues. eLife 12. 10.7554/eLife.91979.3.

15. Mansfield, S.M., and Kino, G.S. (1990). Solid immersion microscope. Applied Physics Letters 57, 2615–2616. 10.1063/1.103828.

16. Qian Wu, Ghislain, L.P., and Elings, V.B. (2000). Imaging with solid immersion lenses, spatial resolution, and applications. Proc. IEEE 88, 1491–1498. 10.1109/5.883320.

17. Barnes, W.L., Björk, G., Gérard, J.M., Jonsson, P., Wasey, J.A.E., Worthing, P.T., and Zwiller, V. (2002). Solid-state single photon sources: light collection strategies. Eur. Phys. J. D 18, 197–210. 10.1140/epjd/e20020024.

18. Yoshita, M., Baba, M., Koshiba, S., Sakaki, H., and Akiyama, H. (1998). Solid-immersion photoluminescence microscopy of carrier diffusion and drift in facet-growth GaAs quantum wells. Applied Physics Letters 73, 2965–2967. 10.1063/1.122645.

19. Wang, L., Bateman, B., Zanetti-Domingues, L.C., Moores, A.N., Astbury, S., Spindloe, C., Darrow, M.C., Romano, M., Needham, S.R., Beis, K., et al. (2019). Solid immersion microscopy images cells under cryogenic conditions with 12 nm resolution. Commun Biol 2, 74. 10.1038/s42003-019-0317-6.

20. Matheson, A.M.M., Chua, N.J., and Tosches, M.A. (2025). Iberian ribbed newts. Current Biology 35, R49–R51. 10.1016/j.cub.2024.11.062.

21. Chung, K., Wallace, J., Kim, S.-Y., Kalyanasundaram, S., Andalman, A.S., Davidson, T.J., Mirzabekov, J.J., Zalocusky, K.A., Mattis, J., Denisin, A.K., et al. (2013). Structural and molecular interrogation of intact biological systems. Nature 497, 332–337. 10.1038/nature12107.

22. Tomer, R., Ye, L., Hsueh, B., and Deisseroth, K. (2014). Advanced CLARITY for rapid and high-resolution imaging of intact tissues. Nature Protocols 9, 1682–1697. 10.1038/nprot.2014.123.

23. Renier, N., Wu, Z., Simon, D.J., Yang, J., Ariel, P., and Tessier-Lavigne, M. (2014). iDISCO: A Simple, Rapid Method to Immunolabel Large Tissue Samples for Volume Imaging. Cell 159, 896– 910. 10.1016/j.cell.2014.10.010.

24. Klingberg, A., Hasenberg, A., Ludwig-Portugall, I., Medyukhina, A., Männ, L., Brenzel, A., Engel, D.R., Figge, M.T., Kurts, C., and Gunzer, M. (2017). Fully Automated Evaluation of Total Glomerular Number and Capillary Tuft Size in Nephritic Kidneys Using Lightsheet Microscopy. J Am Soc Nephrol 28, 452–459. 10.1681/ASN.2016020232.

25. Wang, Q., Ding, S.-L., Li, Y., Royall, J., Feng, D., Lesnar, P., Graddis, N., Naeemi, M., Facer, B., Ho, A., et al. (2020). The Allen Mouse Brain Common Coordinate Framework: A 3D Reference Atlas. Cell 181, 936-953.e20. 10.1016/j.cell.2020.04.007.

26. Guttikonda, S.R., Sikkema, L., Tchieu, J., Saurat, N., Walsh, R.M., Harschnitz, O., Ciceri, G., Sneeboer, M., Mazutis, L., Setty, M., et al. (2021). Fully defined human pluripotent stem cell-derived microglia and tri-culture system model C3 production in Alzheimer’s disease. Nat Neurosci 24, 343– 354. 10.1038/s41593-020-00796-z.

27. Liu, J.T.C., Glaser, A.K., Poudel, C., and Vaughan, J.C. (2023). Nondestructive 3D Pathology with Light-Sheet Fluorescence Microscopy for Translational Research and Clinical Assays. Annual Rev. Anal. Chem. 16, 231–252. 10.1146/annurev-anchem-091222-092734.

28. Ertürk, A. (2024). Deep 3D histology powered by tissue clearing, omics and AI. Nat Methods 21, 1153–1165. 10.1038/s41592-024-02327-1.

29. Song, A.H., Williams, M., Williamson, D.F.K., Chow, S.S.L., Jaume, G., Gao, G., Zhang, A., Chen, B., Baras, A.S., Serafin, R., et al. (2024). Analysis of 3D pathology samples using weakly supervised AI. Cell 187, 2502-2520.e17. 10.1016/j.cell.2024.03.035.

30. Bishop, K.W., Erion Barner, L.A., Han, Q., Baraznenok, E., Lan, L., Poudel, C., Gao, G., Serafin, R.B., Chow, S.S.L., Glaser, A.K., et al. (2024). An end-to-end workflow for nondestructive 3D pathology. Nat Protoc 19, 1122–1148. 10.1038/s41596-023-00934-4.

31. Serafin, R., Xie, W., Glaser, A.K., and Liu, J.T.C. (2020). FalseColor-Python: A rapid intensity-leveling and digital-staining package for fluorescence-based slide-free digital pathology. PLoS One 15, e0233198. 10.1371/journal.pone.0233198.

32. Woych, J., Ortega Gurrola, A., Deryckere, A., Jaeger, E.C.B., Gumnit, E., Merello, G., Gu, J., Joven Araus, A., Leigh, N.D., Yun, M., et al. (2022). Cell-type profiling in salamanders identifies innovations in vertebrate forebrain evolution. Science 377, eabp9186. 10.1126/science.abp9186.

33. Ouellette, N., Recknagel, A., Cao, K., Baka, J., Chandrashekar, J., and Logsdon, M. (2023). Whole Mouse Brain Delipidation, Immunolabeling, and Expansion Microscopy.

34. Steib, E., Vagena-Pantoula, C., and Vermot, J. (2023). TissUExM protocol for ultrastructure expansion microscopy of zebrafish larvae and mouse embryos. STAR Protocols 4, 102257. 10.1016/j.xpro.2023.102257.

35. Asano, S.M., Gao, R., Wassie, A.T., Tillberg, P.W., Chen, F., and Boyden, E.S. (2018). Expansion Microscopy: Protocols for Imaging Proteins and RNA in Cells and Tissues. Curr Protoc Cell Biol 80, e56. 10.1002/cpcb.56.

36. Guenthner, C.J., Miyamichi, K., Yang, H.H., Heller, H.C., and Luo, L. (2013). Permanent Genetic Access to Transiently Active Neurons via TRAP: Targeted Recombination in Active Populations. Neuron 78, 773–784. 10.1016/j.neuron.2013.03.025.

37. Susaki, E.A., Tainaka, K., Perrin, D., Yukinaga, H., Kuno, A., and Ueda, H.R. (2015). Advanced CUBIC protocols for whole-brain and whole-body clearing and imaging. Nat Protoc 10, 1709–1727. 10.1038/nprot.2015.085.

38. Kenney, J.W., Steadman, P.E., Young, O., Shi, M.T., Polanco, M., Dubaishi, S., Covert, K., Mueller, T., and Frankland, P.W. (2021). A 3D adult zebrafish brain atlas (AZBA) for the digital age. eLife 10, e69988. 10.7554/eLife.69988.

39. Rajput, N., Parikh, K., Squires, A., Fields, K.K., Wong, M., Kanani, D., and Kenney, J.W. (2025). Whole-Brain Mapping in Adult Zebrafish and Identification of the Functional Brain Network Underlying the Novel Tank Test. eNeuro 12. 10.1523/ENEURO.0382-24.2025.

40. Lopes Da Costa, B., Helms, K.M., Theodore, K., Tsai, Y.-T., Caruso, S.M., Liu, S., Lima De Carvalho, J.R., Nolan, N.D., Tahir, S., Makinson, C.D., et al. (2025). Prime editing for the investigation of aberrant splicing defect associated with a pathogenic PRPH2 variant. Molecular Therapy Nucleic Acids 36, 102740. 10.1016/j.omtn.2025.102740.

41. Qi, Y., Yu, T., Xu, J., Wan, P., Ma, Y., Zhu, J., Li, Y., Gong, H., Luo, Q., and Zhu, D. (2019). FDISCO: Advanced solvent-based clearing method for imaging whole organs. Science Advances 5, eaau8355. 10.1126/sciadv.aau8355.

42. Schindelin, J., Arganda-Carreras, I., Frise, E., Kaynig, V., Longair, M., Pietzsch, T., Preibisch, S., Rueden, C., Saalfeld, S., Schmid, B., et al. (2012). Fiji: an open-source platform for biological-image analysis. Nat Methods 9, 676–682. 10.1038/nmeth.2019.

43. Rueden, C.T., Schindelin, J., Hiner, M.C., DeZonia, B.E., Walter, A.E., Arena, E.T., and Eliceiri, K.W. (2017). ImageJ2: ImageJ for the next generation of scientific image data. BMC Bioinformatics 18, 529. 10.1186/s12859-017-1934-z.

44. Schneider, C.A., Rasband, W.S., and Eliceiri, K.W. (2012). NIH Image to ImageJ: 25 years of image analysis. Nat Methods 9, 671–675. 10.1038/nmeth.2089.

