## Supplementary Info and Figure for "Hybrid Solid-Liquid Optics Enable Scalable, High-Resolution, Multi-Immersion Light-Sheet Microscopy"

### Supplementary material

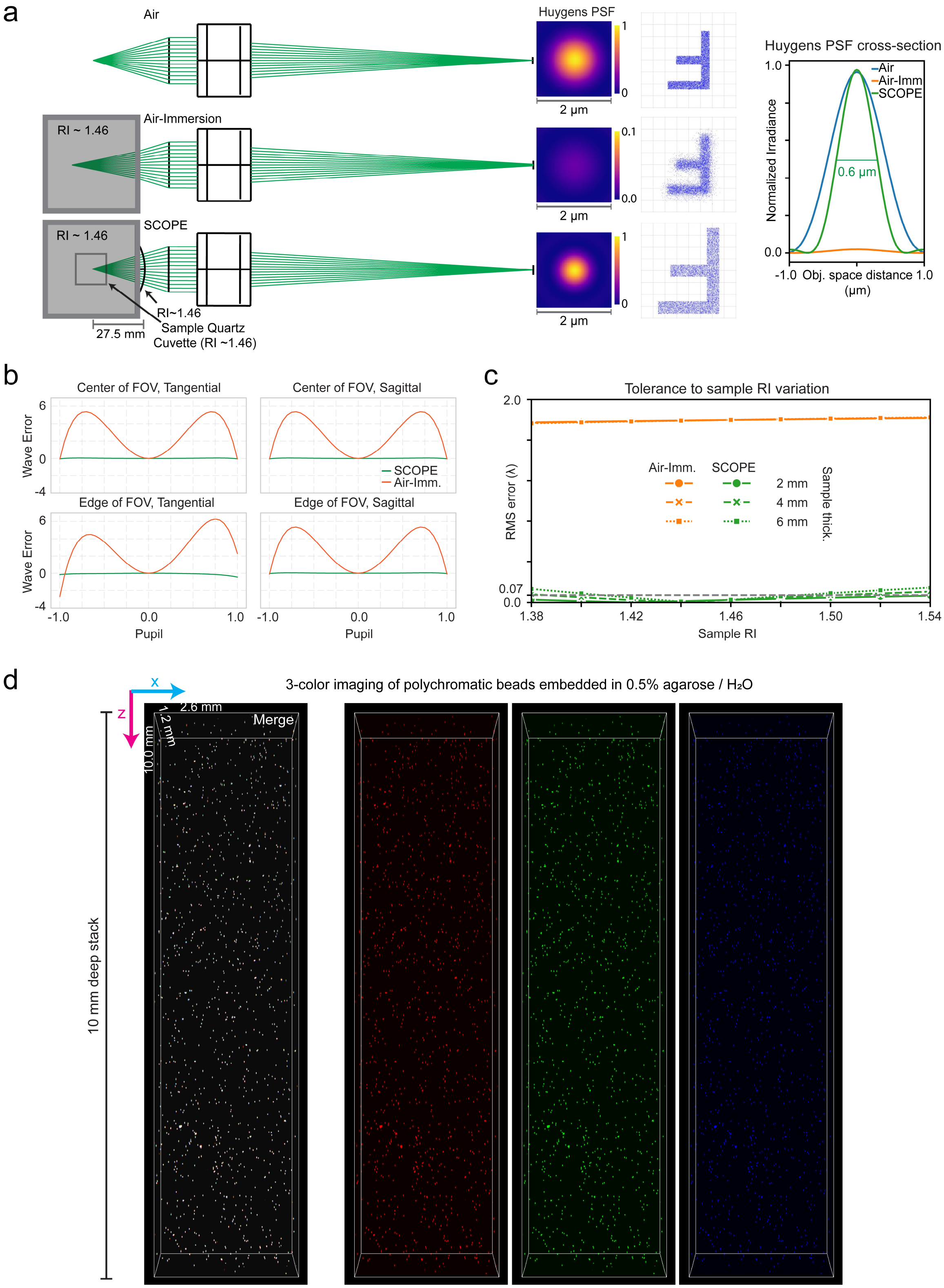


**Supplementary Figure 1 | Deep, aberration-free multicolor imaging with SCOPE across a large field of view and refractive index mismatch. (a)** Zemax optical simulations comparing detection using an air objective in air, in immersion media (standard configuration), and with the SCOPE imaging device. The radius of curvature of the truncated hemispherical (plano-convex; R=27.5 mm) lens used matches the practical configuration employed for all biological imaging experiments. Left to right: optical ray traces, estimated Huygens point-spread functions (PSFs), simulated image formation, and quantification of normalized irradiance profiles of the Huygens PSFs. (**b**) Wavefront error analysis (optical path difference) across sagittal and tangential planes at the center and edge of the field of view, demonstrating aberration correction by SCOPE. (**c**) Root-mean-square (RMS) wavefront error as a function of sample refractive index and thickness. **(d)** Three-dimensional light-sheet imaging of polychromatic fluorescent beads embedded in 0.5% agarose/$H_2O$ (RI ~ 1.33) using the SCOPE imaging device (system RI ~ 1.46). Despite the large refractive-index mismatch between the aqueous sample (~1.33) and the hybrid solid-liquid immersion system (~1.46), SCOPE achieves aberration-free, chromatically aligned imaging across the full sample depth (~1 cm demonstrated) and width (~2.6 mm demonstrated). Left to right: merged multichannel image and individual fluorescence channels (red, green, and blue).

**Supplementary Videos**:

**Supplementary Video 1 | High-resolution volumetric imaging of expanded *Thy1-GFP* mouse brain section using SCOPE.** Flythrough of a sub-volume from a ~4x expanded *Thy1-GFP* mouse brain section imaged with the SCOPE-enabled pLSM/SLICE system with a 10x/0.28 NA/34 mm WD air objective. The bounding box measures 3.41 mm × 2.14 mm × 0.6 mm. Submicron resolution reveals detailed neuronal morphology, including dendritic spines, across large tissue volumes. The dataset represents a small region of interest (ROI) from the full dataset shown in **Figure 3**.

**Supplementary Video 2 | Whole-brain imaging of the dopaminergic system in an expanded salamander brain using SCOPE.** Volumetric rendering of a ~4x expanded adult salamander brain immunolabeled for tyrosine hydroxylase (TH) and imaged with the SCOPE-enabled pLSM/SLICE system with a 10x/0.28 NA/34 mm WD air objective. The bounding box measures 11.84 mm x 8.05 mm x 6.0 mm. The dataset shows fine neuronal processes and features across the entire intact brain. Dataset corresponds to **Figure 4a**.

**Supplementary Video 3 | Visualization of a 200 µm sagittal optical section from the whole salamander brain dataset.** High-magnification view from the same dataset as Supplementary Video 2, showing TH expression across different parts of individual neurons. Dataset corresponds to Figure **4b–c**.

**Supplementary Video 4 | High-resolution whole-brain imaging of a CUBIC-R–cleared fosTRAP2-tdTomato mouse brain.** Volumetric rendering of an intact mouse brain cleared with CUBIC-R and imaged using SCOPE-enabled pLSM/SLICE system with a 10x/0.28 NA/34 mm WD air objective. *tdTomato*$^+$ neurons are clearly resolved throughout the brain. The bounding box measures 21.13 mm x 16.98 mm x 9.55 mm. The dataset is compatible with registration to standard atlases and downstream quantitative analyses, as shown in **Figure 5a**.

**Supplementary Video 5 | High-resolution imaging of an iDISCO-cleared *A. mexicanus* (cavefish) brain labeled with TO-PRO-3.** 3D light-sheet imaging of the nuclear-labeled cavefish brain cleared in organic solvent. SCOPE enables uniform nuclear resolution across the entire brain despite the high refractive index of the clearing medium. Imaging was performed using a 10x/0.28 NA/34 mm WD air objective. The bounding box measures 7.23 mm x 3.19 mm x 2.55 mm. Dataset corresponds to **Figure 5b**.

**Supplementary Video 6 | High-throughput, high-resolution 3D imaging of human iPSC-derived cortical organoids co-cultured with microglia.** Volumetric rendering of day-94 human iPSC-derived cortical organoids co-cultured with microglia and immunolabeled for IBA1. The dataset reveals heterogeneous microglial morphologies - including amoeboid, elongated, and ramified forms - distributed throughout the organoid volume. Imaging was performed using the SCOPE-enabled pLSM/SLICE system with a 10x/0.28 NA/34 mm WD air objective. Dataset corresponds to **Figure 5c**.

**Supplementary Video 7 | Large-volume 3D histopathology of human breast tissue using SCOPE.** Volumetric rendering with maximum-intensity projection of an intact human breast tissue sample imaged using the SCOPE-enabled pLSM/SLICE system with a 10x/0.28 NA/34 mm WD air objective, following fluorescent H&E analog labeling (TO-PRO-3 for nuclei and eosin for cytoplasm), ECi clearing, and false H&E-like coloring. The dataset reveals terminal duct–lobular units (TDLUs) and their tree-like mesoscopic organization at nuclear resolution. Dataset corresponds to **Figure 6a**.

**Supplementary Video 8 | Detailed volumetric rendering of human breast tissue architecture.** Minimum-intensity-projection volumetric rendering of the same dataset shown in Supplementary Video 7, highlighting the organization of TDLUs in tree-like structures. The video demonstrates high-resolution imaging across centimeter-scale tissue volumes. Dataset corresponds to **Figure 6a**.

**Supplementary Video 9 | 3D histopathology of invasive carcinoma in human breast tissue.** Volumetric rendering of a cancerous human breast tissue sample showing invasive carcinoma with fibrotic focus features. The dataset reveals the volumetric architecture of cancerous tissue with subcellular detail, enabling virtual sectioning and 3D histopathological assessment. The dimensions of the full cylindrical volume shown are 1.82 mm in diameter and 1 mm in height. Imaging was performed using the the SCOPE-enabled pLSM/SLICE system with a 10x/0.28 NA/34 mm WD air objective. Dataset corresponds to **Figure 6b**.
